# The dynamic upper limit of human lifespan

**DOI:** 10.1101/124800

**Authors:** Saul Newman, Easteal Simon

**Affiliations:** John Curtin School of Medical Research, Australian National University, Acton, ACT 2601, Australia

## Abstract

We respond to claims by Dong *et al*. that human lifespan is limited below 125 years. Using the log-linear increase in mortality rates with age to predict the upper limits of human survival we find, in contrast to Dong *et al*., that the limit to human lifespan is historically flexible and increasing. This discrepancy can be explained by Dong *et al.’s* use of data with variable sample sizes, age-biased rounding errors, and log(0) instead of log(1) values in linear regressions. Addressing these issues eliminates the proposed 125-year upper limit to human lifespan.

## Main text

Recent findings by Dong *et al*.^1^ suggested fixed upper limits to the human life span. Using the same data, we replicated their analysis to obtain an entirely different result: the upper limit of human life is rapidly increasing.

Mortality rates double with age in human populations (Fig. 1a-b). Log-linear models fit to this rate-increase closely approximate the observed age-specific probability of death^2^. These models also provide a simple method of predicting upper limits to human life span that is independent of population size.

**Figure 1.**
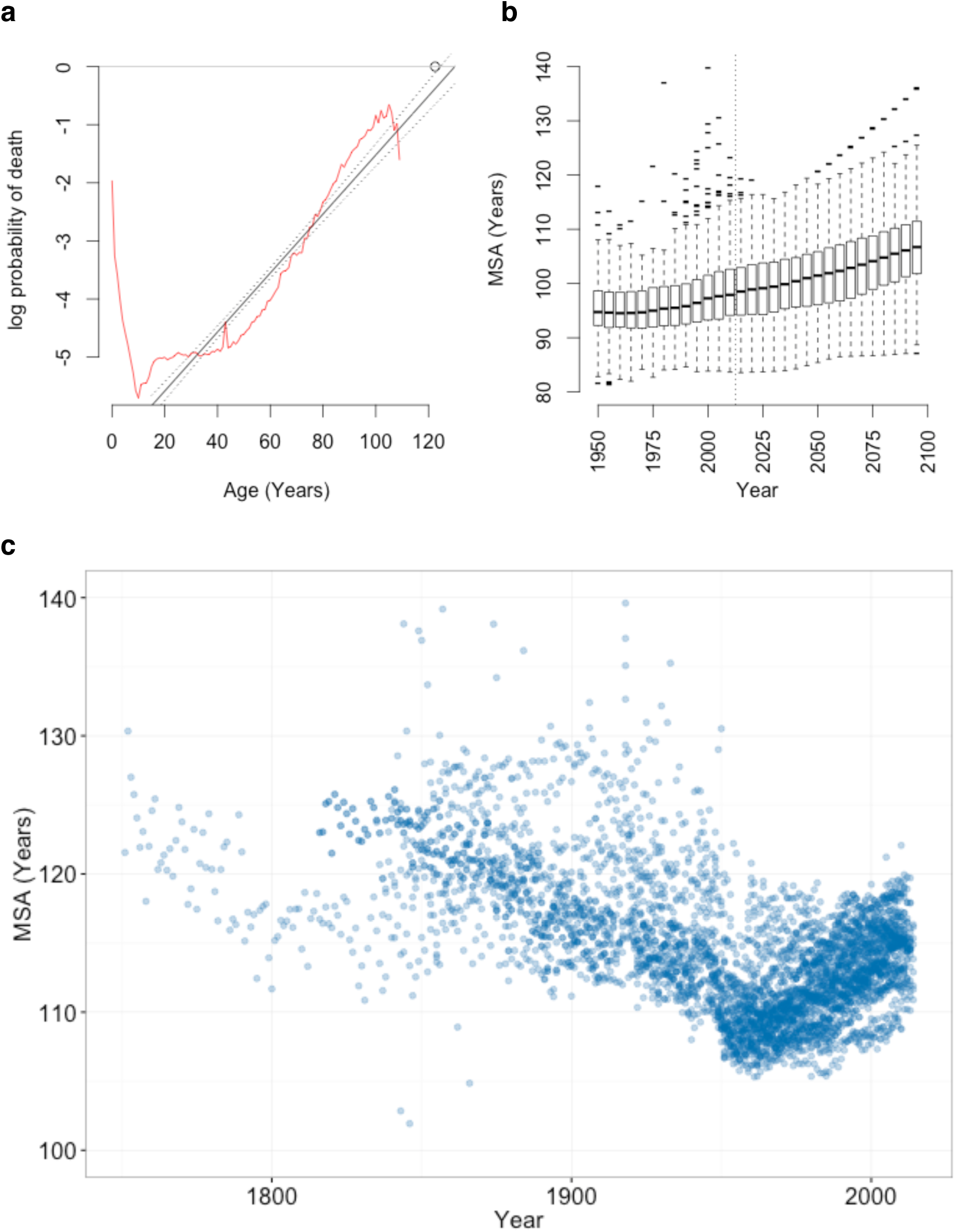
Observed and projected variation in the maximum survivable age (MSA).

**a**, In humans, the probability of death *q* at age *x* (*q*_*x*_; red line) increases at an approximately log-linear rate with age (black lines; 95% CI), shown here for the birth cohort of Jeanne Clament (d.122.5 years; circle). Projection of this log-linear increase to log(*q*) = 0 provides the MSA, the upper limit of human survival, shown here for **b**, observed and projected global populations^4^ and **c**, 40 historic populations^3^ 1751–2014.

Here, we fit log-linear models to age-specific mortality rates from the Human Mortality Database^3^ (HMD) data used by Dong *et al*.^1^ and predict the age at which the probability of death intercepts one. This maximum survivable age (MSA) provides a simple, conservative estimate of the upper limit of human life (Fig. 1c).

Log-linear models closely approximate the observed probability of death in HMD populations for both period and cohort life tables (median R^2^ = 0.99; 4501 population-years). These models predict an MSA exceeding 125 years within observed historic periods (Fig. 2b-c; SI).

**Figure 2.**
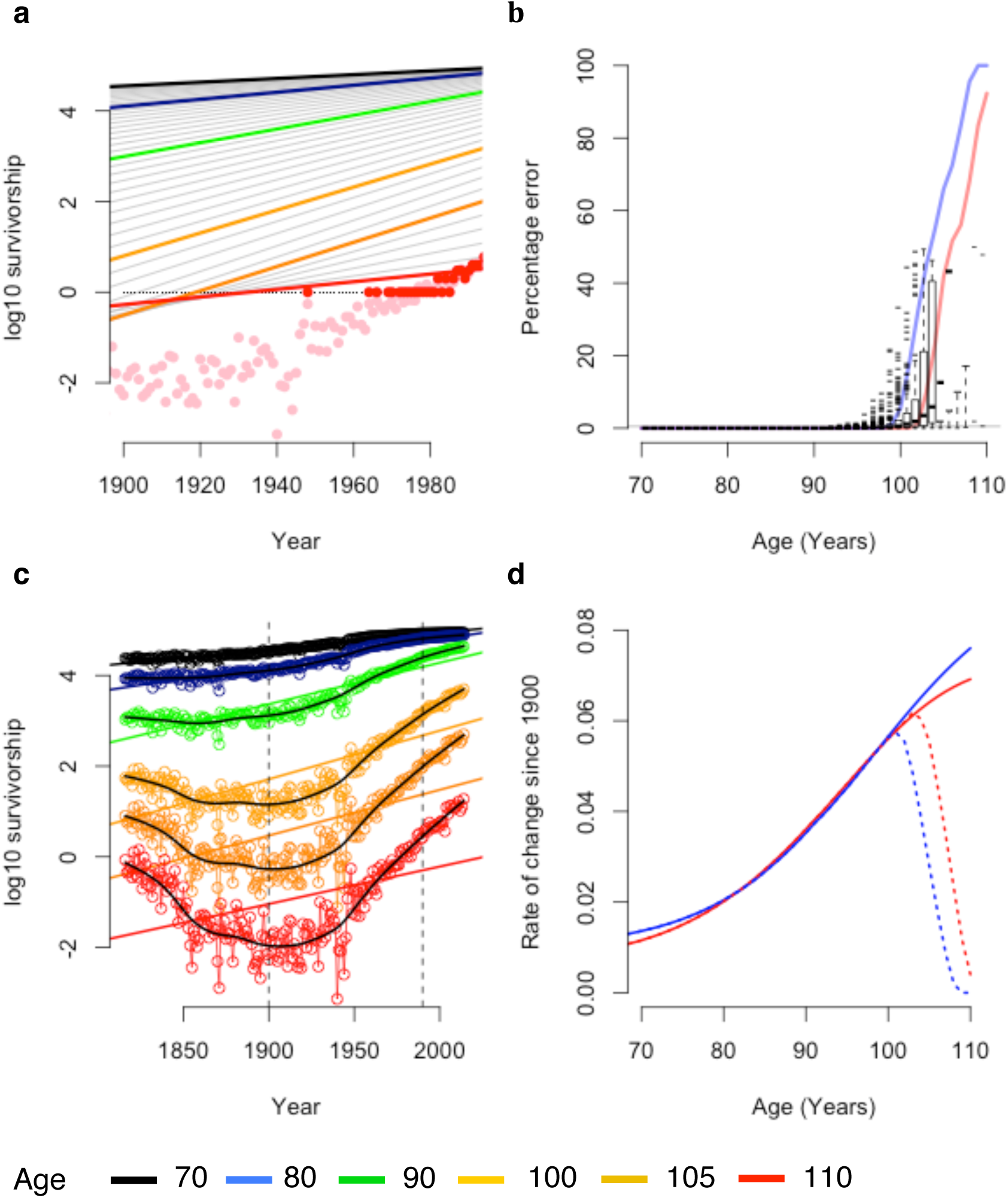
Rate of change in late-life survival for the French population 1816-2014.

**a**, Figure modified after Dong *et al*. Fig. 1b, showing rounded survival data (red points), rounded survival data with log(0)=log(l) (black points), the resulting linear regression in Dong *et al*. (solid red line) and observed survival data (pink points), **b**, Rounding errors in survival data (box-whisker plots; 95% Cl) and the proportion of survival data rounded to zero in males (blue line) and females (red line), **c**, Survival data from **a** with rounding errors removed, showing variation outside the 1900-1990 period (vertical dotted lines), **d**, The rate of change in late-life mortality since 1900 with (dotted lines) and without (solid lines) rounding errors (after Dong *et al*. Fig. 1c).

Furthermore, period data indicate that MSA is steadily increasing from a historic low c.1956 (Fig. 2b-c) and that the maximum reported age at death (MRAD) is expected to rise over the next century. This result is supported by trends in global mortality data from the United Nations^4^ sampled across 194 nations (Fig. 1b).

This analysis provides an estimate of human lifespan limits that is conservatively low. Log-linear mortality models assume no late-life deceleration in mortality rates^5^, which, if present, would increase the upper limits of human lifespan^6^. In addition, these models are fit to population rates and cannot provide an estimate of individual variation in the rate of mortality acceleration.

These proposed limits and are discrepant with Dong *et al*.^1^. Dong *et al*. conclude that the MRAD is limited to 125 years in humans^1^ and that lifespan increases above age 110 are highly unlikely, due to the reduced rate of increase in life expectancy at advanced ages.

To resolve this discrepancy we replicated Dong *et al*.’s^1^ analysis using identical data (SI). Replicating these findings requires the inclusion of rounding errors, treating zero-rounded values as log(1) and the incorrect pooling of populations.

The HMD data provide both the age-specific probability of survival (*q*_*x*_) and the survival rates of a hypothetical cohort of 100,000 individuals (*l*_*x*_). However, *l*_*x*_ survival rates are rounded off to the nearest integer value.

The magnitude and frequency of *l*_*x*_ rounding errors increases as the probability of survival approaches 1 in 100,000. These rounding errors mask variation in survival rates at advanced ages: over half of *l*_*x*_ survival data are rounded to zero above age 90 (Fig. 2b).

Dong *et al*. appear to have used these rounded-off survival data in their models^1^ and incorrectly treated log(0) values as log(1) in log-linear regressions (Fig. 2a-d; SI).

These errors have considerable impact. Re-calculating cohort survival from raw data or excluding zero-rounded figures eliminates the proposed decline in old-age survival gains (Fig. 2d; SI).

Likewise, recalculating these data removed their proposed limits to the age of greatest survival gain (SI), which in 15% of cases were the result of the artificial 110-year age limit placed on HMD data^7^.

We also found that variation in the probability of death was masked by date censoring^1^. Major non-linear shifts in old-age survival occur outside the 1900-1990 period used by Dong *et al*. (Fig. 2c). Why these data were excluded from this regression, but included elsewhere, is unclear.

Evidence based on observed survival above age 110 appears to support a late-life deceleration in survival gains^1^. For the period 1960-2005, Dong *et al*. present data^1^ from 4 of the 15 countries in the International Database on Longevity^8^ (IDL). In their pooled sample of these countries, there is a non-significant (p=0.3) reduction in MRAD between 1995 and 2006 (Fig. 3a).

**Figure 3.**
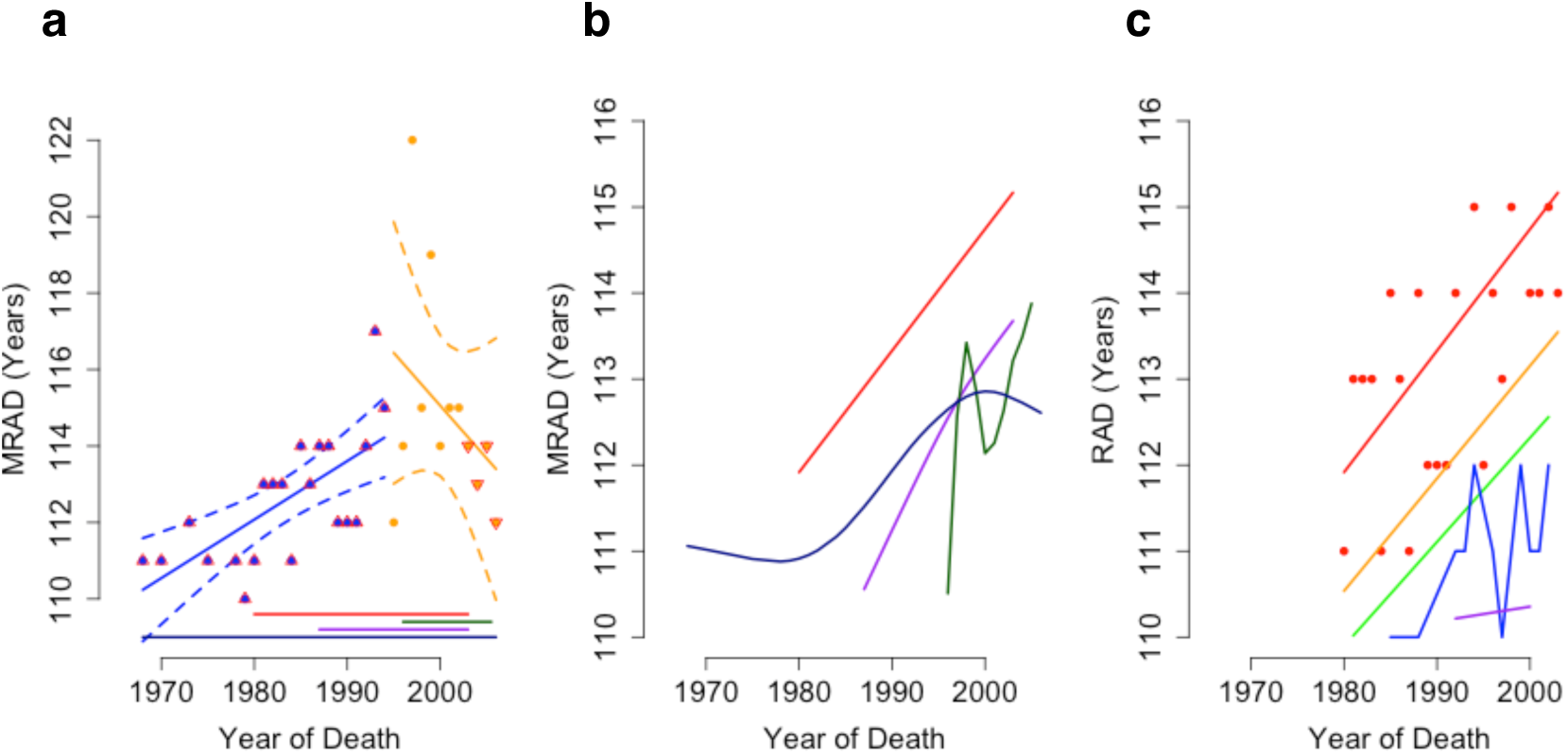
Maximum reported age at death of supercentenarians.

**a**, Reproduction of Dong *et al*. Figure 2a, including 95% CI for increasing (p<0.0001) and falling (p=0.3) maximum recorded age at death (MRAD), showing data biased by the addition and removal (up and down arrows) of populations. **b**, Locally weighted smoothed splines of MRAD in Japan (green), the USA (red), the UK (dark blue) and France (purple). **c**, Locally weighted trends of MRAD in the USA across the oldest 5 reported ages at death (red, orange, green, blue and purple lines show rank 1-5 respectively).

The declining MRAD reported by Dong *et al*.^1^ arise from the use of falling sample sizes. Of the validated^9^ supercentenarians alive in 2007, 62% lived in France and the USA. However, these countries are not surveyed^8^ by the IDL after 2003 (Fig. 3a). The proposed post-1995 decline in MRAD results from this dramatic fall in sample size.

Viewed individually, all four countries have an upward trend in the mean reported age at death (RAD; Fig. 3b) of supercentenarians (SI) and the top 5 ranked RADs (Fig. 3c). All four countries achieved record lifespans since 1995, as did 80% of the countries in the IDL. Without the pooling of IDL data used by Dong *et al*. there is no evidence for a plateau in late-life survival gains.

We attempted to reproduce Dong *et al*.’s supporting analysis of Gerontology Research Group^9^ (GRG) records. The text and figure S6 do not match annual MRAD records from 1972 as stated^1^. However, they do match deaths of world’s oldest person titleholders from 1955 (GRG table C, revision 9) with all deaths in May and June removed (SI).

Actual MRAD data from the GRG support a significant decline in the top-ranked age at death since 1995 (r = −0.47; p = 0.03, MSE = 3.2). However, this trend is not significant if only Jeanne Clement is removed (p = 0.9). Linear models fit to lower-ranked RADs have an order of magnitude better fit, and all indicate an increase in maximum lifespan since 1995 (N= 64; SI).

Collectively these data indicate an ongoing rebound of upper lifespan limits since 1950, with a progressive increase in the theoretical and observed upper limit of human life. Given historical flexibility in lifespan limits and the possibility of late-life mortality deceleration in humans^10^, these models should, however, be treated with caution.

A claim might be made for a general, higher 130-year bound to the human lifespan. However, an even higher limit is possible and should not be ruled out simply because it exceeds observed historical limits.

## Author contributions

S.J.N. wrote the analysis and code, and reproduced Dong *et al.’s* analysis. S.J.N. and S.E. developed the analysis, methods and statistical design, and co-wrote the manuscript.

## Corresponding Author

S.E.

## Methods

Life table data were downloaded from the United Nations^4^ (UN) and the Human Mortality Database^11^ (HMD) and lifespan records from the International Database on Longevity^12^ (IDL) and the Gerontology Research Group^9^ (GRG).

Least squared linear models were fit to life table data on the log-transformed age-specific probability of death (*q_x_*), and projected to *q_x_*=1 to predict the maximum survivable age in each population (Fig. 1b-c; SI). Maximum lifespan within GRG and IDL data was annually aggregated and fit by locally weighted smoothed splines^13^ (Fig. 3b,c).

We reproduced the analysis of Dong *et al*. in R version^14^ 3.2.1 (SI).

## Data Availability

The authors declare that all data are available within the paper and its supplementary files.

## Code availability

All code is freely available from the authors on request.

## References

1. Dong, X., Milholland, B. & Vijg, J. Evidence for a limit to human lifespan. Nature 538, 257–259 (2016).

2. Finch, C. E. Variations in senescence and longevity include the possibility of negligible senescence. J Gerontol A Biol Sci Med Sci 53, B235–9 (1998).

3. Human Mortality Database. (2013). at <www.mortality.org>

4. United Nations. World Population Prospects: The 2015 Revision. United Nations Econ. Soc. Aff. XXXIII, 1–66 (2015).

5. Rose, M. R. et al. Evolution of late-life mortality in Drosophila melanogaster. Evolution 56, 1982–1991 (2002).

6. Thatcher, A. R. The long-term pattern of adult mortality and the highest attained age. J. R. Stat. Soc. 162, 5–43 (1999).

7. Wilmoth, J. R., Andreev, K., Jdanov, D. & Glei, D. A. Methods Protocol for the Human Mortality Database Version 5.0. 83 (2007). at <http://www.mortality.org/Public/Docs/MethodsProtocol.pdf>

8. Maier H, Gampe J, Jeune B, R.J. Supercentenarians and V. J. Supercentenarians. Demogr. Res. Monogr. 7, (2010).

9. Gerontology Research Group. at <http://www.grg.org/SC/SCindex.html>

10. Kannisto, V., Lauritsen, J., Thatcher, A. R. & Vaupel, J. W. Reductions in Mortality at Advanced Ages: Several Decades of Evidence from 27 Countries. Popul. Dev. Rev. 20, 793–810 (1994).

11. Max Planck Institute for Demographic Research. Human Mortality Database. University of California, Berkeley and INED, Paris at <http://www.mortality.org/>

12. Robine, J. & Planck, M. IDL, the International Database on Longevity. Gerontology 1–30 (2005).

13. Cleveland, W. S. LOWESS: A program for smoothing scatterplots by robust locally weighted regression. Am. Stat. 35, 54 (1981).

14. R Core Development Team. R: A language and environment for statistical computing. R Foundation for Statistical Computing, Vienna, Austria. {ISBN} 3-900051-07-0 (2012).

